# Cross-species discovery of large structural variants reveals distinct evolutionary dynamics in primates

**DOI:** 10.64898/2026.08.28.747789

**Authors:** Feifei Zhou, Junmin Han, Shilong Zhang, Jiadong Lin, Evan E. Eichler, Yafei Mao

## Abstract

Large-scale structural variants (SVs) are major drivers of genome evolution and disease susceptibility, shaping lineage-specific gene content, genome architecture, and phenotypic diversity. Yet, their systematic detection across divergent species has been hindered by complex rearrangements and reference bias, limiting our understanding of their evolutionary impact. Here we present LGvar, a divergence-aware tool for identifying SVs in population-scale and cross-species genome assemblies, enabling up to a 7-fold increase in the detection of large deletions, insertions, inversions and complex rearrangements (≥10 kbp). Application of LGvar uncovers 112 previously undetected large SVs overlooked by conventional methods and quantifies the effect of evolutionary divergence on SV discovery across primate euchromatic genomes, revealing an approximately 30% overall decline in detection sensitivity with increasing phylogenetic distance. Mapping SVs from comparable syntenic regions onto the primate phylogeny shows that large deletions (≥10 kbp) accumulate approximately two-fold faster than insertions and eight-fold faster than inversions, in contrast to the dynamics of smaller SVs (<10 kbp). Integrating structurally divergent regions with the synteny region analysis reveals over 606 previously unreported gene gains and losses in great apes and refined the totals to 134 human-lineage-specific and 51 great-ape-lineage-specific protein coding genes. These results suggest that large SVs follow distinct evolutionary trajectories, with potential selective constraints shaping genome structure. Our study establishes a generalizable strategy for cross-species SV discovery and provides a high-resolution view of how large SVs contribute to genome evolution in primates.

## Introduction

Understanding genetic variation among individuals and across species is fundamental to uncovering the molecular basis of phenotypic diversity, evolutionary adaptation, speciation and human diseases^1–6^. Among all forms of genomic variation, structural variants (SVs)— including insertions, deletions, inversions, duplications, and translocations—represent one of the most impactful yet technically challenging classes to identify. SVs can reshape gene structures, alter regulatory landscapes, and mediate genome rearrangements^7–9^. In both humans and other organisms, they are strongly associated with phenotypic innovation and disease susceptibility^10–15^. For example, inversions in deer mice influence tail length variation and local adaptation^11^; large-scale rearrangements in primate genomes modulate gene expression across different cell types^12^; and recurrent microdeletions such as 16p11.2 and 2q13 are strongly implicated in neurodevelopmental disorders^13,14^.

Early genomic studies at the turn of the 21st century could only detect megabase-scale copy number variations (CNVs) using array-based technologies^16–19^ . The advent of short-read sequencing enabled detection of insertions and deletions but still left over 70% of SVs unresolved due to limitations in read length and mappability, particularly within repetitive or segmental duplication (SD) regions^20,21^. WGS-based deletion callers vary substantially in sensitivity and precision across coverage levels and variant-size ranges, with no single method performing best under all conditions^22^. The development of long-read sequencing technologies (e.g., PacBio HiFi and Oxford Nanopore) has revolutionized SV discovery, revealing over 25,000 variants per individual genome when compared to the human reference^23–25^. However, accurately resolving large and complex rearrangements such as duplications, translocations, or nested inversions remains a major challenge. The problem becomes even more pronounced in cross-species comparisons, where genome divergence and reference bias further complicate SV detection.

To address these challenges, a growing number of SV detection tools have been developed, generally falling into two methodological classes: read-based and assembly-based approaches^26,27^. Tools such as Sniffles^28^, PBSV (https://github.com/PacificBiosciences/pbsv), SVIM^29^, NanoSV^30^ and cuteSV^31^ leverage raw read alignments to detect breakpoints and are well-suited for low-coverage population studies. In contrast, assembly-based methods such as PAV^32,33^ and SVIM-asm^34^ provide higher sensitivity and precision by directly comparing assembled haplotypes, but require more computational resources and high-quality assemblies. More recently, machine learning–based frameworks have been introduced, transforming sequence alignment features into image-like representations for variant calling (e.g., DeepVariant^35^, SVision^36^, SVision-pro^37^). Despite these advances, most current methods have been optimized for human or population-scale studies and show limited performance in identifying large-scale rearrangements or cross-species genomic variation. SyRI^38^, a widely used cross-species comparative genomics tool, excels in detecting large interspecies structural rearrangements but fails to capture smaller, nested, or sequence-level SVs and other genetic variants.

With the rapid generation of high-quality telomere-to-telomere (T2T) assemblies^39,40^ from large human consortia (HPRC^41^, HGSVC^32^, CPC^42^, APG^43^) and from non-human species (VGP^44^, PGC^3^, EBP^45^), there is a growing need for new computational frameworks that can systematically and accurately identify all classes of genomic variation—both within populations and across species.

Here, we present LGvar, a tool designed to enable population and species-level SV discovery from long-read assemblies. LGvar enables comprehensive detection of both small and large-scale genomic variation, including inversions and complex rearrangements. Applying this tool to recent primate T2T assemblies, we systematically reconstruct the evolution of large SVs (>=10 kbp), revealing both reference bias and distinct evolutionary patterns among these variants. Beyond improving large SV detection sensitivity and precision, LGvar provides a robust foundation for comparative genomics and evolutionary analyses across diverse taxa.

## Results

### LGvar overview and performance benchmarking across simulated, population, and cross-species data

Large-scale genomic variations—such as inversions, duplications, and structurally divergent/complex regions (SDRs^12^)—represent some of the most challenging classes of SVs to resolve using read/assembly-based approaches. These difficulties arise primarily from noisy whole-genome alignments, nested inversions, and rearrangements that combine multiple event types. To overcome these limitations, we developed LGvar, a tool designed for robust discovery of complex genomic variation across individuals and species. LGvar identifies genomic variants from whole-genome alignments through a multi-step workflow consisting of alignment filtering, variant discovery, refinement, genotyping, and output generation (Fig. 1; Supplementary Figure 1 and 2; See Methods).

**Figure 1.**
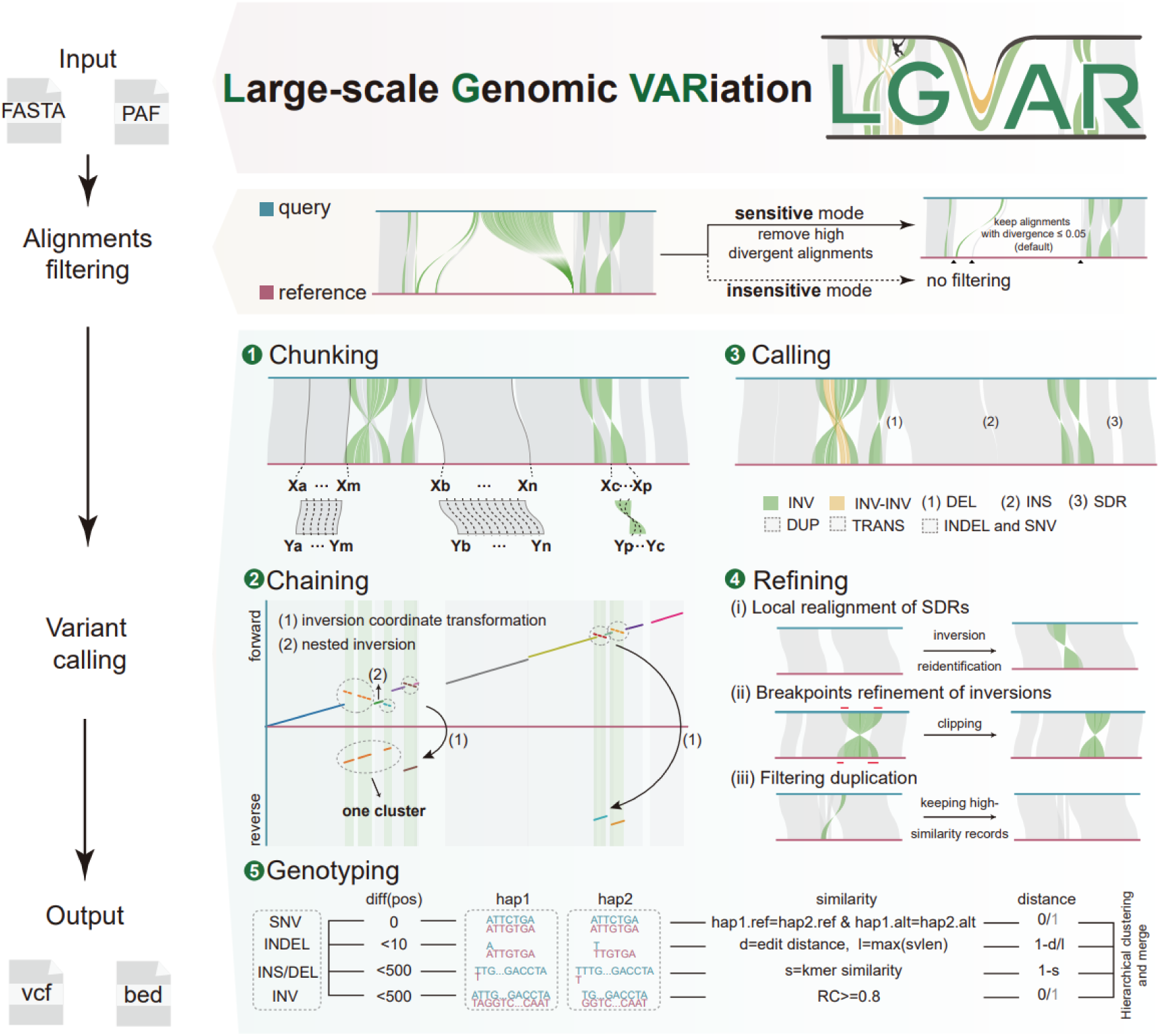
The LGvar workflow for structural variant discovery in genome assemblies. LGvar proceeds in three stages. Input and alignment filtering: assembled genomes (FASTA) or pre-computed alignments (PAF) are accepted as input; divergent alignments are removed in sensitive mode or retained in insensitive mode. Variant calling: five modules operate in series — chunking partitions query (blue) and reference (red) alignments into segments; chaining clusters segments and resolves nested inversions; calling identifies inversions (INV), nested inversions (INV–INV), segmental duplication–associated regions (SDR), insertions (INS), deletions (DEL), duplications (DUP), translocations (TRANS), indels and single-nucleotide variants (SNV); refining polishes candidate calls by local realignment of SDRs, breakpoint adjustment and duplication filtering; and genotyping assigns genotypes by hierarchical clustering on position, haplotype, sequence similarity and breakpoint distance. Output: variants are exported in VCF and BED format.

**Figure 2.**
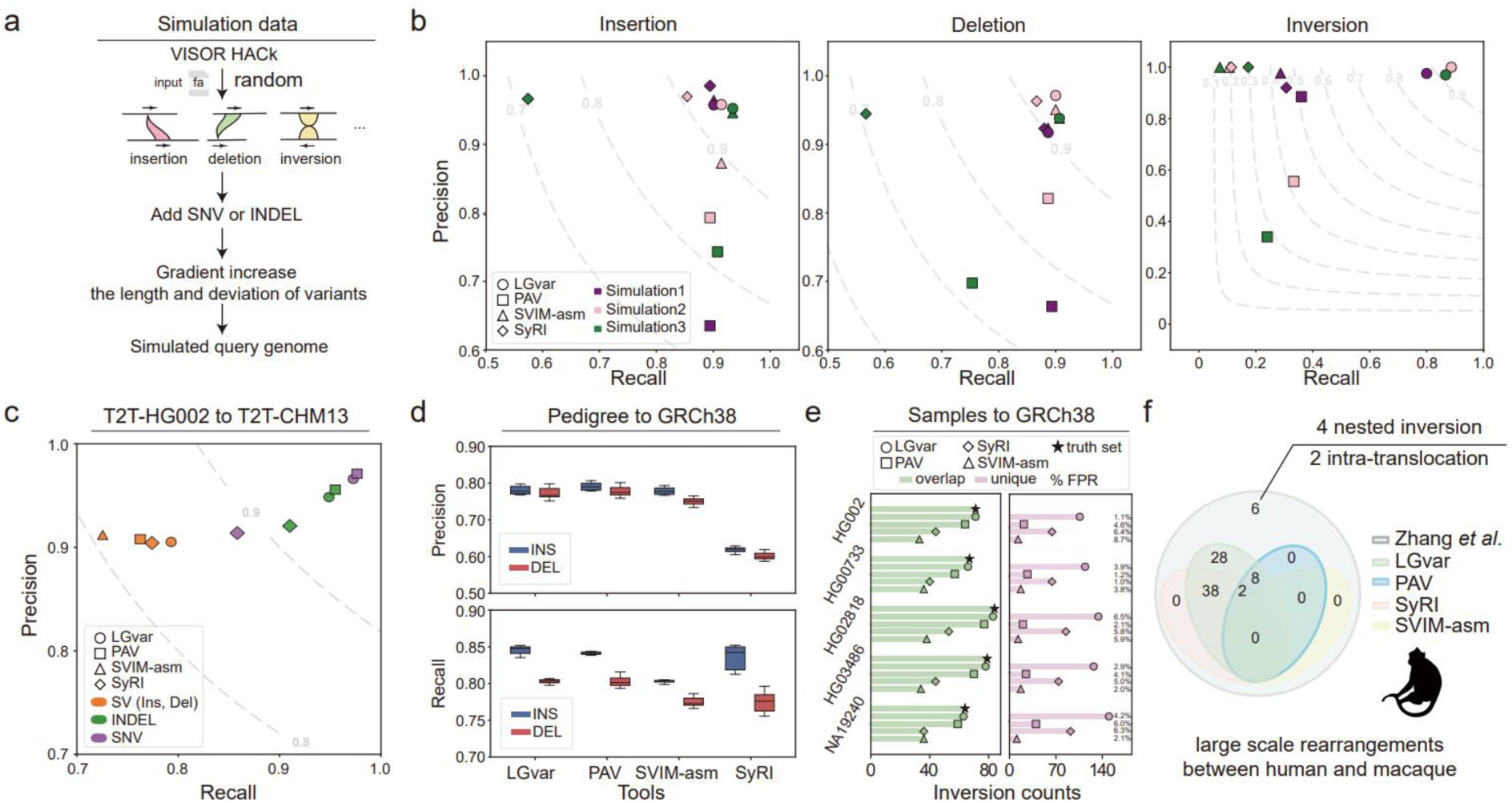
Benchmarking LGvar against other structural variant callers using simulated data and empirical primate genomes. (a) Schematic overview of the simulated data generation pipeline (Methods). (b) Performance comparison of multiple SV callers on simulated datasets containing insertions, deletions, and inversions. (c) SV detection performance on the T2T-HG002 genome aligned to T2T-CHM13, evaluated using LGvar, PAV, SVIM-asm, and SyRI. (d) Precision and recall for insertion (blue) and deletion (red) detection across seven pedigree genomes aligned to GRCh38. Each marker (shape and color) represents a unique combination of genome and SV caller. (e) Inversion detection performance across five genomes relative to GRCh38, assessed using LGvar, PAV, SVIM-asm, and SyRI. The left panel shows the number of overlapping inversions between call sets and the truth set, while the right panel shows inversions uniquely identified by each tool; false positive rates (FPRs) are indicated. (f) Benchmarking against previously reported large-scale rearrangements identified in comparisons between human and crab-eating macaque genomes. Venn diagrams show the number of known events detected by each tool.

First, LGvar filters highly divergent alignments to reduce the influence of poorly aligned or highly diverged genomic regions. By default, the sensitive mode retains alignments with sequence divergence below 0.05, while the insensitive mode allows users to bypass this filtering step.

Second, LGvar performs variant discovery using five sequential modules. In the chunking module, whole-genome alignments between the query and reference genomes are partitioned into syntenic segments according to chromosomal coordinates. Because whole-genome alignments often prioritize global consistency over local precision, LGvar further divides syntenic and inverted regions into non-overlapping 5-kbp windows based on aligned coordinate distances to improve local resolution. In the chaining module, chunked windows are clustered separately according to forward and reverse orientations. This step includes coordinate transformation for inversions and resolution of nested inversions. The windows are then grouped using the DBSCAN^46^ algorithm to filter out small or noisy alignments and connect contiguous segments into coherent alignment blocks representing conserved or rearranged regions.

The calling module performs initial identification of diverse variant types, including inversions (INV), nested inversions (INV-INV), structurally divergent regions (SDRs), insertions (INS), deletions (DEL), duplications (DUP), translocations (TRANS), INDELs, and SNVs. The refining module further polishes raw calls by locally realigning SDRs to identify hidden inversions, refining inversion breakpoints using clipping information, and filtering duplications based on sequence similarity.

Finally, the genotyping module assigns genotypes by integrating positional differences, haplotype consistency, sequence similarity, and genomic distance, followed by hierarchical clustering and merging of redundant or overlapping calls. In the final output stage, LGvar exports the validated genomic variants in standard VCF and BED formats for downstream analysis.

To rigorously evaluate the performance of LGvar relative to established assembly-based SV detection tools—PAV^32,33^, SVIM-asm^34^, SyRI^38^—we conducted a comprehensive benchmarking analysis using both simulated and real datasets representing a range of genomic complexity and evolutionary divergence (Supplementary Table 1, 2).

We first assessed tool performance using simulated genomes derived from human chromosome 1. Using the *HACk* module of VISOR^47^, we introduced random sets of insertions, deletions, and inversions at three levels of increasing sequence divergence, each defined by progressively larger mean variant lengths and standard deviations (Fig. 2a; Supplementary Figure 3 and 4; see Methods). Across all sequence divergence levels, LGvar demonstrated consistently high accuracy for insertions and deletions, while showing a striking advantage in inversion detection—a class of large-scale variation that remains challenging for most assembly-based methods. LGvar achieved a precision near 1.0 and a recall above 0.8, in contrast to competing tools, all of which exhibited inversion recall below 0.4 (Fig. 2b; Supplementary Table 3). This pronounced difference underscores LGvar’s robustness in resolving complex and large-scale structural rearrangements.

**Figure 3.**
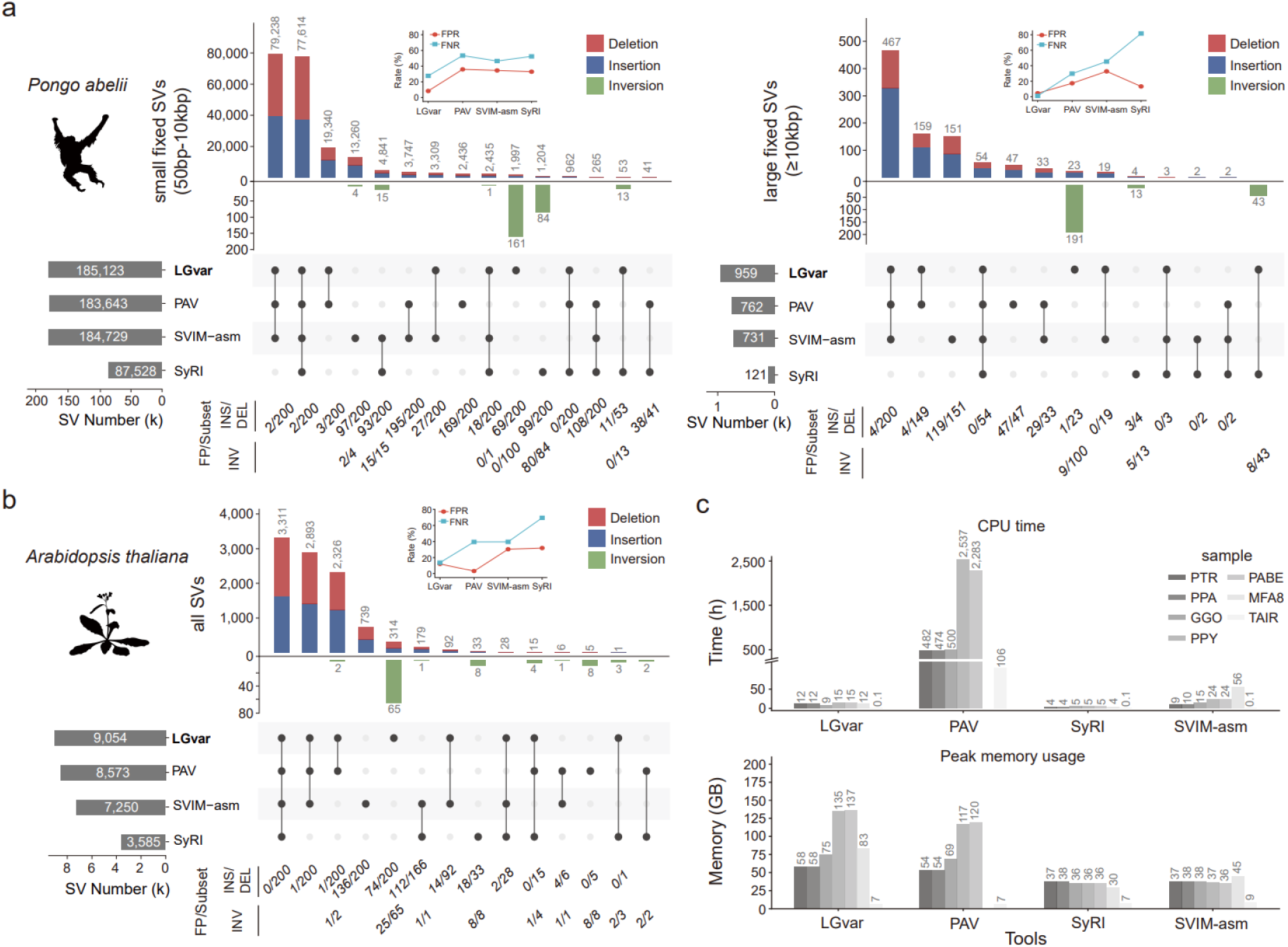
Structural variant identification across primate species and in *Arabidopsis thaliana* genomes. (a) UpSet plots showing the numbers of SVs identified by each caller in the *Pongo abelii* genome aligned to the T2T-CHM13 reference. SV types are color-coded: deletions (red), insertions (blue), and inversions (green). False-positive (FP) counts for each caller are shown below the UpSet plots, with corresponding false-positive rates (FPRs) and false-negative rates (FNRs) displayed on the line chart. Small and large fixed SVs are shown in the left and right panels, respectively. (b) UpSet plot summarizing SVs identified from a comparison between *Arabidopsis thaliana* accessions Col-0 and Ler. False-positive (FP) counts for each caller are shown below the UpSet plots, with corresponding false-positive rates (FPRs) and false-negative rates (FNRs) displayed on the line chart. (c) Computational resource usage, including CPU time and peak memory, for the analyses shown in a and b. The dashed box indicates missing data for PAV due to runtime errors.

**Figure 4.**
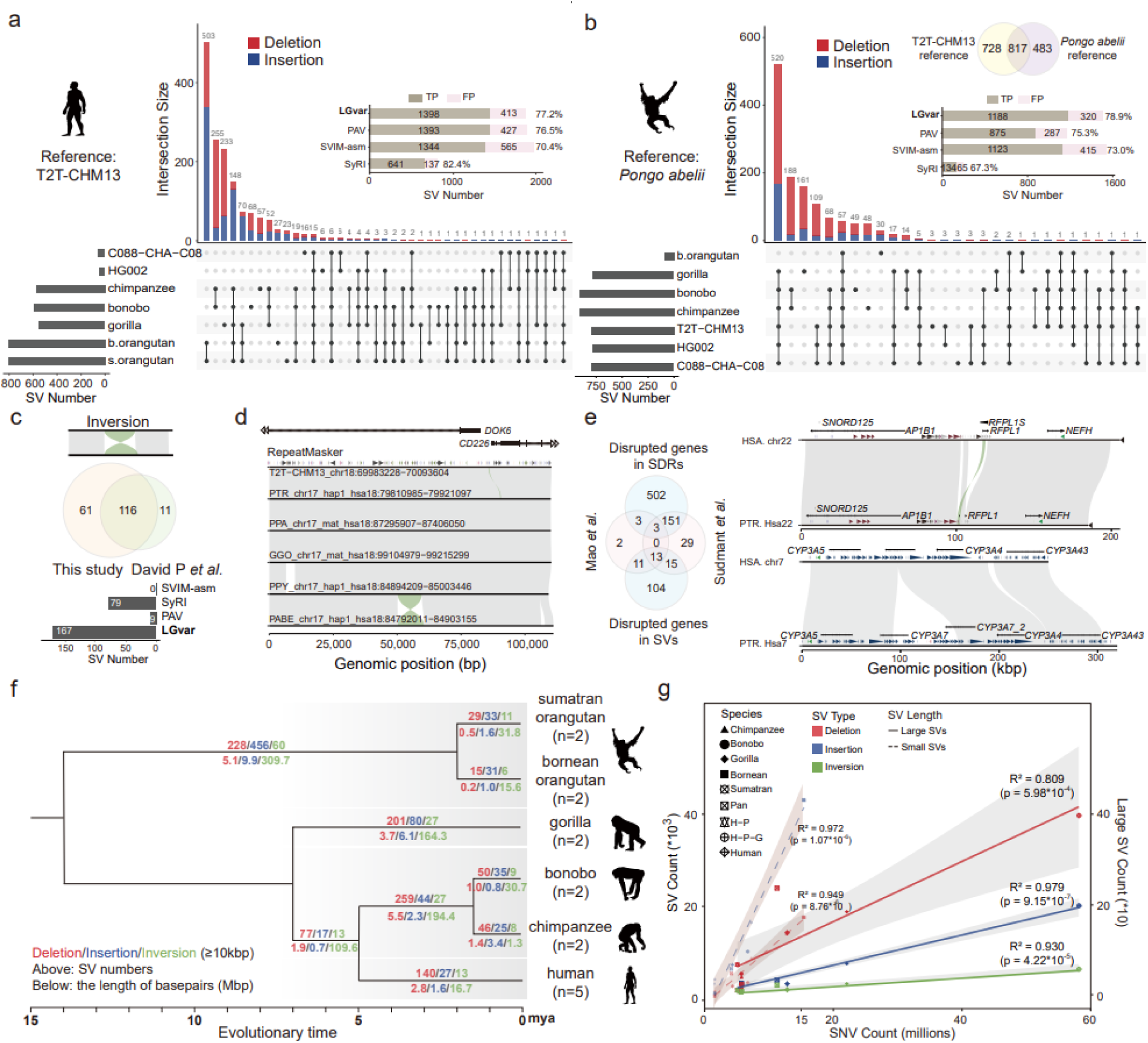
Large-scale structural variation across primate genomes. (a, b) UpSet plots showing the distribution of insertions and deletions across great ape lineages. The total number of SVs detected by each caller and the corresponding false positives (FPs) are indicated. T2T-CHM13 (a) and the Sumatran orangutan genome (b) were used as reference assemblies. The venn diagram shows the overlap between the two mapping directions. (c) Benchmarking of lineage-specific inversions using the dataset from Porubsky et al., *Cell*, 2022. (d) Examples of newly identified lineage-specific inversions. (e) Genes disrupted by large-scale SVs benchmarked against the dataset from Mao *et al.*, *Cell*, 2024 and Sudmant *et al.*, *Genome Res*, 2013. The Venn diagram summarizes disrupted genes shared between datasets. The two newly reported examples are shown in the right panel, including *Pan*-lineage specific *RFPL1* gene loss and human-lineage specific *CYP3A7_2* gene loss. (f) Numbers of large-scale fixed structural variants—deletions (red), insertions (blue), and inversions (green)—mapped onto each branch of the primate phylogeny. For each branch, the number of events is shown above the line and the cumulative size (Mbp) below. Numbers assigned to ancestral branches denote lineage-specific SVs (for example, the *Pan* lineage shared by bonobo and chimpanzee). (g) Correlation between the number of fixed large-scale structural variants and the accumulation of single-nucleotide variants across lineages, shown separately for deletions (red), insertions (blue), and inversions (green). Correlations for small- and large-scale SVs are indicated by dashed and solid lines, respectively.

We next benchmarked LGvar using high-quality human genome assemblies. Our initial evaluation used the T2T-HG002 assembly from the Human Q100 Project^48^, which provides a well-curated variant benchmark. Both LGvar and PAV achieved high accuracy and substantially outperformed SyRI and SVIM-asm. While PAV yielded marginally higher F1 scores for SNV and INDEL detection (difference <0.01), LGvar achieved slightly higher F1 scores for SV detection (difference <0.02) (Fig. 2c; Supplementary Table 4). In addition, the LGvar also show higher performance on F1 on INDEL detection on SVA elements, suggesting LGvar have better ability to deal with INDELs in complex TE (Supplementary Figure 5).

We next assessed performance in a population context using seven assemblies from the CEPH-1463 pedigree generated by the Platinum Pedigree Consortium^49^ (Fig. 2d). Consistent with the single-genome benchmark, LGvar and PAV again showed comparable accuracy and markedly outperformed SyRI and SVIM-asm. On average, LGvar exhibited a precision difference of <0.01 lower than PAV, while achieving a recall <0.01 higher than PAV (Fig. 2d; Supplementary Figure 6-7; Supplementary Table 5). Overall, LGvar maintained competitive performance across SNVs, INDELs, and SVs. Notably, these benchmark datasets do not include validated inversions, precluding evaluation of large-scale rearrangements in this setting.

To directly evaluate inversion detection, we analyzed five human genome assemblies^50^ with empirically validated inversion call sets. LGvar recovered 98-100% of known inversions, whereas alternative methods detected only 48-90% (Fig. 2e). In addition to the validated events, LGvar identified 106-140 additional inversion candidates per genome, representing up to a four-fold increase in discovery yield relative to existing callers (Supplementary Table 6; Supplementary Figure 8). Manual curation of these additional calls indicated that LGvar achieves approximately 1.2-fold lower false discovery rates compared with competing methods, demonstrating improved sensitivity without compromising precision in inversion detection. In addition, we identified 85 inversions in HG002 relative to the T2T-CHM13 reference assembly, comprising 16 homozygous and 69 heterozygous events (Supplementary Table 7). Approximately 22% and 50% of inversion breakpoints coincided with TEs and SDs, respectively, consistent with a role for repeat-mediated rearrangement in their formation^51^ (Supplementary Figure 9).

We next assessed LGvar’s ability to identify large-scale genomic rearrangements in a validated cross-species T2T comparison dataset (T2T-CHM13 human vs. T2T-MFA8 macaque), which contains 82 manually curated SVs exceeding 100 kbp in size^52^. LGvar successfully recovered 92.7% (76/82) of reported rearrangements with default parameters, compared with 48.8% (40/82) for SyRI, 12.2% (10/82) for PAV, and 0% (0/82) for SVIM-asm (Fig. 2f; Supplementary Table 8). Among the 6 undetected variants in LGvar, four were nested inversions, and two corresponded to translocation.

Together, these analyses demonstrate that LGvar achieves performance comparable to, or exceeding, state-of-the-art assembly-based SV detection tools across both simulated and empirical datasets. Its advantages are most pronounced in identifying inversions and other large-scale genomic rearrangements, where traditional approaches often fail.

### Species-scale genetic variation discovery comparison

Because cross-species benchmarking datasets with validated truth sets remain limited, we extended our evaluation to comparative genome analyses to assess LGvar’s ability to resolve evolutionary rearrangements and divergence in genome architecture at the species scale.

We applied LGvar alongside three existing methods to identify evolutionary SVs from cross-species genome comparisons using five telomere-to-telomere (T2T) nonhuman primate (NHP) assemblies^40^ (Supplementary Table 1, 2). Using the human-orangutan comparison as a representative example, LGvar detected approximately 0.8%-112% more small SVs (insertions and deletions, 50 bp-10 kbp) than the other methods, while achieving the lowest false positive rates (FPRs) and false negative rates (FNRs) (Fig. 3a). For large SVs (≥10 kbp), LGvar identified 0.26 to 7-fold more variants relative to alternative tools (Fig. 3a; Supplementary Table 9). These results indicate improved sensitivity while maintaining high specificity, particularly for complex evolutionary rearrangements.

We further assessed cross-species performance using a plant dataset comprising *Arabidopsis thaliana* accessions Col-0 and Ler^38^. LGvar identified 0.06 to 1.5-fold more variants relative to alternative tools, again with the lowest FPRs and FNRs (Fig. 3b; Supplementary Table 10).

We next examined the calls that were private to individual tools. Approximately 40% of these were attributable to differences in breakpoint representation that persisted through merging, rather than to differences in detection; the remainder reflected true differences in caller sensitivity and specificity. Manual inspection of the latter set showed that LGvar had the lowest FPRs in both the human-orangutan and the *Arabidopsis thaliana* comparisons.

We additionally compared computational performance across methods. PAV required the longest runtime, consuming on average 85-fold more CPU time than LGvar (Fig. 3c; Supplementary Table 11). On average, LGvar showed a peak memory usage comparable to that of PAV. SyRI showed the shortest runtime and lowest memory usage; however, it detected 112%-693% fewer SVs than LGvar and exhibited 9-25% higher FPRs. While PAV performs robustly in human SV detection, its performance and computational efficiency appear less suitable for cross-species analyses involving extensive genome rearrangements.

Together, these cross-species evaluations demonstrate that LGvar is well suited for large-scale comparative genomics, enabling sensitive and reliable detection of SVs across animal and plant genomes.

### Large-scale lineage-specific SVs in primates

The above analyses demonstrate that LGvar is a robust tool for comparative genomics and large-scale SV characterization. Leveraging this capability and integrating complementary evidence from additional tools, we next addressed a fundamental evolutionary question: what are the lineage-specific large SVs (≥10 kbp) in primates, and at what rate do they arise during evolution? In parallel, we systematically evaluated an important technical concern that has not been comprehensively addressed—namely, the impact of reference bias on cross-species SV discovery.

In addition to mapping NHP genomes to the human T2T-CHM13 reference, we performed reciprocal analyses by mapping human and other NHP genomes to the orangutan (*Pongo abelii*) reference genome. We first integrated SV calls from all tools by svpop^53^ and manually curated each event (total 2,811). Among the evaluated methods, LGvar exhibited the lowest FPR (∼21%), whereas SyRI showed the highest FPR (33%).

These reciprocal analyses revealed a pronounced reference bias. Approximately 60% of true-positive SVs were detected in only one of the two mapping directions (Fig. 4a-b; Supplementary Table 12-14), largely reflecting increasing genetic divergence between reference and query genomes. For example, when using the human genome as the reference, 16 additional true SVs were detected in Bornean orangutan versus human comparisons; however, when using Sumatran orangutan as the reference, 19 additional true SVs were identified for the same species pair. These results demonstrate that reference choice substantially influences SV discovery in comparisons involving large evolutionary distances.

To mitigate reference bias, we integrated the validated SVs from both mapping strategies to reconstruct the most comprehensive set of primate lineage-specific large SVs, including insertions, deletions, and inversions. Previous studies have reported large-scale lineage-specific inversions in primates using different genome assemblies and analytical frameworks^54^. Benchmarking our inversion call set against these studies showed that 91.3% (116/127) of previously reported inversions were recovered (Fig. 4c). Of these, 15.5% (18/116) had inconsistent lineages (Supplementary Figure 10). In addition, we identified 61 novel lineage-specific inversions, 97% (59/61) of which were detected by LGvar, with 57% (35/61) uniquely identified by LGvar (Fig. 4d; Supplementary Table 15-16; Supplementary Figure 11-12). The eleven inversions not recovered in our analysis were classified as low-confidence events in the prior study^54^, underscoring the improved precision and completeness of our call set.

Finally, we projected the 1,967 large-scale lineage-specific SVs (≥10 kbp) detected in the syntenic/comparable genomics regions onto the primate phylogeny and found that 49.5 Mbp and 874.1 Mbp of genomic sequence were affected by lineage-specific insertions/deletions and inversions, respectively (Fig. 4f, Supplementary Figure 13). Notably, we found that deletions occurred at approximately twice the frequency of insertions among large SVs, a pattern that contrasts sharply with previous observations for smaller lineage-specific SVs (<10 kbp)^12^. This discrepancy likely reflects the distinct mutational mechanisms underlying different size classes of SVs, as small lineage-specific variants are predominantly driven by transposable element (TE) insertions like *Alu/L1* insertions (Supplementary Figure 16). Furthermore, we observed that large lineage-specific SVs are correlated with SNV divergence across lineages and accumulate at rates approximately eight-fold and two-fold slower than those of small lineage-specific insertions and deletions (Fig. 4g; Supplementary Figure 17).

Next, we identified 1,041 SDRs across primate genomes, collectively encompassing 229.1 Mbp of the human genome. Owing to the complex rearrangements within these regions, SDRs can rarely be assigned unambiguously to individual evolutionary lineages. We therefore focused instead on the lineage-specific gene gains and losses associated with SDRs, which can be treated as discrete characters and mapped onto the primate phylogeny.

Together, lineage-specific large-scale SVs and SDRs affected 143 and 659 protein-coding genes, respectively, 134 of which were human-specific (Supplementary Figure 14). Our callset captured 86.3% (196/227) of the genes identified in previous studies, while also uncovering 606 novel disrupted genes not previously reported^12,55^ (Fig. 4e, Supplementary Table 17-21, Methods). Among the 134 human-lineage genes, four are absent from the previously reported human-specific catalogue^56^: *AGAP4* (vesicular trafficking), *CLEC18C* (innate immunity), *FOXO3B* (metabolism and stress response) and *SPACA5B* (reproductive development).

Functional enrichment analysis revealed that these entire genes disrupted by SVs and SDRs are enriched in pathways related to chemosensory perception (*p* = 4.03e-18), metabolism (*p* = 0.00261), and other biological processes (Supplementary Table 22; Supplementary Figure 15), suggesting potential roles in lineage-specific traits and adaptive evolution. For example, *RFPL1*, a gene involved in cell-cycle regulation, is specifically depleted in the *Pan* lineage. Given its reported role in neural cell-cycle control^57^, this loss may be associated with aspects of *Pan* brain evolution (Fig. 4e). In addition, we identified a tandem duplication of the metabolic gene *CYP3A7* in other NHPs that is absent in humans, indicating a human-specific structural configuration that may have functional consequences^58^ (Fig. 4e). Furthermore, two inversions occurring at the human-*Pan* ancestral node were found to remodel the untranslated regions (UTRs) of *TOP3A* and *YTHDC1*—critical regulators of DNA surveillance and RNA epitranscriptomic modification, respectively^59,60^. The potential for these SVs to alter the regulatory landscape of such fundamental cellular processes warrants further functional validation to elucidate their role in the divergence of the Hominini.

## Discussion

Accurate characterization of genetic variation within and between populations and species is fundamental to population genetics and comparative genomics^61–63^. Although long-read sequencing and T2T genome assemblies have become increasingly feasible, robust identification of large-scale genomic variation and cross-species genetic differences remains challenging. In this study, we developed LGvar, an assembly-based variant discovery tool that integrates refined alignment and variant calling strategies. LGvar demonstrates consistent and competitive performance for SNVs, INDELs, and simple/small SVs (<10 kbp) in human genomes, while substantially outperforming existing tools for inversion and large SV (≥10 kbp) detection in both human and cross-species comparisons.

At the species scale, SV discovery has historically been underexplored, in part because accurate cross-species genome alignment remains technically demanding. Precise alignment is essential for downstream variant calling, particularly for large rearrangements. In LGvar, the chunking and chaining strategy substantially improves alignment resolution, enabling more accurate detection of inversions and other complex SVs/SDRs. The “Chunk and Chain” strategy used in LGvar also reflects an inherent trade-off in whole-genome alignment and structural variant discovery: maintaining global alignment consistency versus maximizing local structural resolution. In LGvar, large alignments are divided into syntenic chunks to reduce the propagation of local alignment errors caused by highly divergent sequences, translocations, or complex rearrangements, and these chunks are subsequently chained to maintain genome-wide collinearity. However, this design may reduce resolution at complex local breakpoints, particularly in repetitive or structurally polymorphic regions where unique anchors are sparse and alignments are ambiguous. To address this issue, LGvar includes a downstream refinement step that performs local realignment in highly polymorphic regions and clipping-based refinement of inversion breakpoints. Nevertheless, breakpoint resolution in highly repetitive or complex regions remains dependent on the quality of the underlying alignment. Future developments could incorporate adaptive chunking, repeat-aware alignment, and other refinement strategies to further improve local variant resolution while preserving global alignment consistency. Notably, SDR detection is therefore sensitive to the choice of DBSCAN parameters, which should be tuned for each genome comparison; 500 kbp performed well empirically across primate genomes (Supplementary Figure 18).

Inversions have long been among the most challenging forms of genetic variation to characterize accurately, particularly when they occur in complex or repetitive genomic regions^50,64^. LGvar improves inversion detection by explicitly handling reverse-oriented alignments through orientation-aware chaining and by using a Chunk-and-Chain strategy to resolve local rearrangement signals that may be obscured in coarse whole-genome alignments. In addition, LGvar re-examines gaps and SDRs through local refinement, enabling the recovery of inversions that may be missed during the initial alignment-based calling process.

Benchmarking against the latest inversion validation datasets highlights that no single method can yet recover all inversion events. Consequently, comprehensive detection of large SVs— especially inversions and complex rearrangements—continues to benefit from integrating multiple tools and manual curation. Thus, a further limitation of this study is that part of the benchmark set relies on manual curation, including visual inspection of whole-genome alignments and dotplots. Although this approach provides high-confidence validation for complex SVs, particularly in repetitive or structurally ambiguous regions, it limits scalability. Future benchmarking efforts, especially for cross-species and non-human genome studies, will require high-quality benchmark sets analogous to GIAB/HG002. However, the construction of such resources requires extensive manual curation, integration of multiple lines of evidence, and substantial computational effort. Therefore, future work should focus on developing more scalable strategies for benchmark-set construction, such as automated candidate extraction, batch visualization of alignment evidence, incorporation of orthogonal sequencing or assembly support, and machine-learning-assisted prioritization or classification. These approaches may reduce the burden of manual inspection while maintaining the reliability of expert-curated benchmark sets.

Primate genomes, built over two decades of high-quality sequencing, provide an unparalleled system to study SVs across species-scales^39,40,65^. Early analyses focused on large CNVs^66,67^, whereas recent T2T assemblies now resolve the full size spectrum within a single framework. What is known of primate SVs nonetheless derives almost entirely from events below 10 kbp, where insertions outnumber deletions two to three-fold^12^. Above 10 kbp the polarity reverses, with deletions exceeding insertions by a comparable margin. The insertion-deletion ratio in syntenic sequence is therefore scale-dependent.

These estimates apply only to sequence that can be aligned and polarized. In SDRs, nested and recurrent rearrangements cannot be resolved into discrete, polarizable events, so we shifted the unit of inference from variants to genes, which remain assignable to lineages where variants do not. On this measure SDRs are gain-biased, duplication-driven gains exceeding losses by ∼2:1. No single directional bias therefore governs SVs in primate genomes. Syntenic sequence is insertion-biased at small scales and deletion-biased at large ones, while SDRs remain a persistent source of gene gain. Analyses confined to syntenic regions capture the conservative majority of the genome and miss the compartment in which most gene innovation occurs.

In addition, fixed inversions also accumulate more slowly than insertions and deletions, likely due to recombination suppression and selective constraints, reflecting an evolutionary strategy to maintain genome integrity. Recurrent inversions, observed in human and macaque populations, appear relatively frequent^51,52,54^, but precise quantification of their rates relative to fixed inversions requires standardized datasets and consistent analytical frameworks.

We also systematically evaluated reference bias in comparative genomics and generated a high-quality curated set of primate SVs. Although reference bias has been widely discussed in evolutionary genomics^68–70^, its specific impact on SV discovery has not been thoroughly quantified. Our results demonstrate that increasing genetic divergence between reference and query genomes reduces alignment precision and consequently diminishes SV detection sensitivity. To mitigate this effect, we recommend reciprocal reference analyses when studying lineage-specific SVs, particularly in comparisons involving long evolutionary distances. Moreover, the accuracy of SV detection depends on the quality of the input assemblies; lower-quality assemblies can lead to increased false-positive rates (Supplementary Figure 19).

Accurate SV characterization in highly repetitive regions remains a major challenge for all genome-wide variation discovery tools. Although assembly-based approaches generally provide improved resolution compared with read-based methods by enabling more continuous sequence comparison and clearer breakpoint-level evidence, complex rearrangements remain difficult to fully resolve even from assembled genomes^71^. Recent analyses of nearly complete human genomes have similarly shown that complex genetic variation remains challenging to characterize at the population scale, particularly in repetitive regions and SDRs^72^. Repetitive genomic regions, including heterochromatin, centromeres, SDs, and long tandem repeat arrays, often generate ambiguous alignments because many repeat copies share extremely high sequence similarity and can vary substantially in copy number and organization among individuals^73^. In these regions, unique anchoring sequences are frequently absent or too sparse, making it difficult to determine the correct placement and orientation of reads or assembly contigs. In addition, repeat expansions, contractions, nested repeats, and structural rearrangements can disrupt collinearity between assemblies and the reference genome, further complicating breakpoint resolution. Therefore, reliable variant calling in heterochromatic and other highly repetitive regions remains limited by the accuracy of the underlying alignments. Importantly, centromeric variation often involves array-level copy-number changes, HOR/monomer composition, repeat homogenization, and internal rearrangements, which may require analytical representations beyond conventional alignment-based SV calls, such as VAMPIRE^74^. Future developments will require repeat-aware alignment algorithms and specialized models adapted to the sequence features of different repeat classes. In future versions, LGvar could incorporate multiple alignment strategies or interface with repeat-specific modules to improve variant characterization in repetitive regions beyond the current implementation. Overall, LGvar advances large-scale and cross-species SV discovery and provides a powerful tool for investigating genome evolution and SVs across diverse taxa.

## Methods

### Workflow of LGvar

#### Two main modules in LGvar

##### Module 1. Chunk and chain genomic segments

Step 1. First, LGvar filters highly divergent alignments to reduce the influence of poorly aligned or highly diverged genomic regions. By default, the sensitive mode retains alignments with sequence divergence below 0.05, while the insensitive mode allows users to bypass this filtering step.

Step 2. Strand orientation: to improve inversion detection and minimize orientation bias, all alignments on the negative strand are transformed into a unified coordinate system. For instance, an alignment originally spanning (X₁, Y₁) to (X₂, Y₂) on the negative strand is converted to (X₁, -Y₁) and (X₂, -Y₂), ensuring uniform coordinate treatment for downstream clustering.

Step 3. Local segmentation and clustering: whole-genome alignments often favor global consistency at the expense of local precision. To enhance local resolution, LGvar divides both syntenic and inverted regions into non-overlapping 5-kbp windows based on distance between aligned coordinates. Then, these windows are clustered using the DBSCAN^46^ (v1.1.12) algorithm, which effectively filters out small or noisy alignments and groups contiguous segments into coherent alignment blocks representing conserved or rearranged regions (Fig. 1).

##### Module 2. Variant discovery

Step 1. Detection of translocations: clusters larger than 1 Mbp are designated as syntenic blocks. LGvar calls a translocation when a non-syntenic cluster lies within the genomic coordinates of a syntenic block but displays a divergent linear alignment trend (Fig. 1). This enables precise identification of rearranged fragments embedded within conserved contexts (Supplementary Figure 1).

Step 2. Inversion identification and refinement: inversions are detected based on negative-strand clusters and their chained positive-strand counterparts. Breakpoints are refined when at least 10% reciprocal overlap exists between syntenic clusters, and adjusted to ensure non-overlapping coordinates. To accommodate complex evolutionary rearrangements, nested inversions (INV-INV) are also identified from hierarchically arranged inverted clusters (Supplementary Figure 2).

Step 3. Discovery of SDRs, insertions, deletions, and duplications: We next examined non-syntenic clusters to identify SDRs and other local SVs. Variants are detected hierarchically: (1) tandem duplications are defined as segments overlapping >50% with two or more syntenic regions. (2) insertions and deletions are inferred based on coordinate discrepancies between reference and query genomes. (3) regions showing multiple overlapped alignments or no alignments are classified as SDRs, representing structurally divergent/complex but unresolvable rearrangements.

Step 4. Reclassification of complex SDRs: to account for inversion signals embedded within SDRs, LGvar reprocesses SDRs iteratively. An SDR is reclassified as an inversion if refined local alignment reveals an inverted orientation of the query segment. This step ensures accurate interpretation of complex inversion-like structures.

Finally, following variant discovery, LGvar performs genotyping by integrating multiple complementary indices, including edit distance, sequence similarity, and reciprocal overlap, to consolidate variant calls across both haplotypes (Fig. 1, see methods). This multi-index integration ensures high-confidence variant assignment while avoiding redundancy between allelic events. The resulting unified call set is then exported in standard VCF and BED formats, enabling seamless downstream analyses such as population-level comparison, cross-species evolutionary inference, and genotype–phenotype association studies.

#### LGvar input and alignment preprocessing

LGvar accepts either assembled genome sequences in FASTA format or whole-genome alignment files in PAF format containing CIGAR strings as input. When FASTA files are provided, whole-genome alignments are generated using minimap2^75^ (v2.30-r1287). When PAF files are provided, the alignment information is directly used in downstream analyses.

To ensure robust and reliable alignments, we first filter out alignments originating from highly complex genomic regions, including centromeres, telomeres, and other regions provided by users. We then extract translocation signals and define one-to-one syntenic blocks between the reference and query genomes (see details in Chaining and clustering section).

#### Chaining and clustering of local alignments

The segmented alignments are subsequently subjected to chaining and clustering, which constitute the core steps of LGvar. Within each syntenic block, alignments are partitioned into non-overlapping 5 kbp segments based on Euclidean distance in the reference-query coordinate space for refining the syntenic blocks with universal criteria.

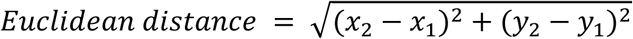

Clustering is performed using the DBSCAN algorithm, with parameters set to an *ε* (epsilon, Eps) of 500 kbp (default), a minimum number of points (MinPts) of 1, and a minimum alignment length threshold of 300 kbp (default) to exclude short or synteny-disrupting alignments (Fig. 1).

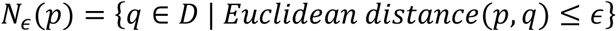

For a given point *p* in a dataset *D*, the *ε*-neighborhood, denoted by *N_ε_*(*p*), is defined as the set of points *q* whose distance to *p* does not exceed a specified radius *ε*.

Alignment segments with Euclidean distances less than or equal to the Eps threshold are chained into the same syntenic cluster. This local alignment refinement strategy substantially improves alignment robustness in highly divergent genomic regions and enhances the accuracy of downstream variant detection.

#### Inversion-aware coordinate transformation

To improve sensitivity for inversion detection, coordinates of inverted alignments are transformed prior to clustering. Specifically, an alignment with original reference and query coordinates (*x*_1_,*y*_1_) and (*x*_2_,*y*_2_) is converted to (*x*_1_,-*y*_1_) and (*x*_2_,-*y*_2_), respectively. This transformation enables inverted segments to be grouped into distinct clusters during the chaining process (Fig. 1), facilitating accurate identification of inversion events.

#### Structural variant detection and classification

Translocation events are identified as alignments that disrupt the continuous synteny clusters. In detail, a translocation is called when a non-syntenic cluster is located within the genomic coordinates of a syntenic block but exhibits a divergent linear trend in the reference-query coordinate space (Fig. 1).

Inversions are detected based on reversed alignment segments and their corresponding chained clusters. Breakpoints are refined when inverted clusters exhibit a reciprocal overlap of at least 0.1 with forward-oriented clusters, after which boundaries are adjusted to eliminate overlapping regions. To accommodate complex evolutionary scenarios in cross-species analyses, LGvar additionally identifies nested inversions (INV-INV) from hierarchically inverted clusters (see details in the section below).

Following the detection of translocations and inversions, LGvar identifies additional variant classes between clusters, including duplications, SDRs, insertions, and deletions. To identify other SVs, LGvar first defines the breakpoints of duplicated regions by detecting overlapping clusters. Alignment records situated between these breakpoints are categorized as duplication events. Then, LGvar identifies SDRs. An SDR is defined as an unaligned gap between two adjacent clusters (e.g., Cluster 1 and Cluster 2) where the genomic distance exceeds 10 kbp. Formally, if Cluster 1 has reference and query coordinates (B, b) at its terminus and Cluster 2 begins at (C, c), an SDR is defined by the intervals (B, C) and (b, c) provided that either |C - B|≥10kbp or |c - b|≥10kbp. Finally, LGvar classifies these SDRs into large insertions or deletions based on a user-defined distance (-d) parameter: Insertion: Defined if the reference gap is negligible (|C - B| < d) while the query gap remains substantial. Deletion: Defined if the query gap is negligible (|c - b| < d) while the reference gap remains substantial.

After inter-cluster variant detection, LGvar further characterized variants within syntenic clusters, including SNVs, INDELs, SDRs, insertions, and deletions by parsing CIGAR strings within alignments (Fig. 1).

#### Iterative refinement of inversion events

Some SDRs may represent small inversions but are not called above. To capture these cases, LGvar iteratively re-evaluates the reference-query alignment. An SDR is reclassified as an inversion if all constituent alignments are inverted or the cumulative aligned length exceeds the threshold of the total SDR length. This refinement step substantially improved the sensitivity and completeness of small-scale inversion detection.

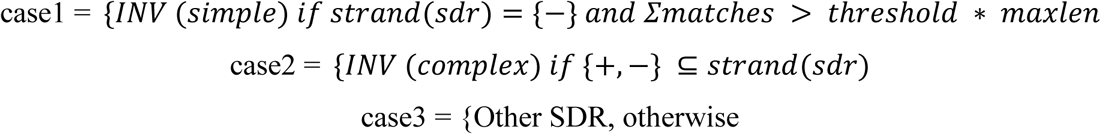

where threshold is calculated as the product of the user-defined fraction parameter and the region length 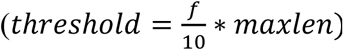, *maxlen* represents the total genomic distance of the SDR, calculated as max (C-B, c-b).

#### Nested inversion events

Nested inversions were identified by a second, orientation-normalized round of clustering. Within each initial alignment cluster, alignments were partitioned by strand into forward (+) and reverse (−) sets. Each forward alignment was then reflected in query coordinates, converting its alignment diagonal from positive to negative slope, and the reflected alignments were pooled with the unmodified reverse-strand alignments for a second clustering pass (DBSCAN). A forward-strand segment was called as an inner inversion when (i) its reflected representation was assigned to the same cluster as reverse-strand alignments and (ii) its reference and query intervals lay within the span of that reverse-strand cluster with flanking inverted sequence on both sides. Candidates were further required to contain at least one original forward-oriented segment, with individual segments contributing ≤50% of the total aligned length and a reference-to-query length ratio of ≤2. Accepted events were reported as nested inversions (INV-INV), with coordinates given for both the outer and the inner interval (Supplementary Fig. 20).

#### Genotyping and output generation

Following variant discovery, LGvar performed genotyping and variant integration through a two-stage clustering approach. First, all variants from both haplotypes were partitioned into groups using a greedy proximity-based algorithm. The grouping distance was adaptively determined by the variant type: a maximum reference distance of 500 bp (*--max_distance*) was applied to SVs, while a smaller distance of 10 bp (*--small_distance*) was used for INDELs. This step ensures that only potentially allelic variants are compared in the subsequent stage.

Second, within each partition group, a similarity-based hierarchical clustering was executed using the complete linkage method. The similarity (s) between any two variants were quantified by integrating specific genomic metrics: base-level edit distance for small INDELs, k-mer Jaccard similarity for large insertions and deletions, and reciprocal overlap (RO) for inversions.

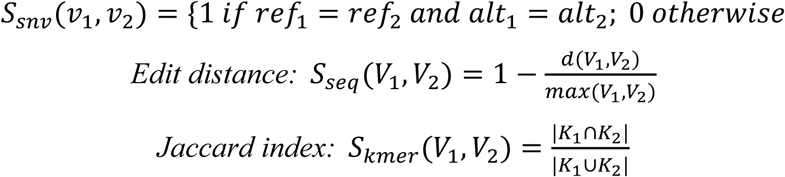

where *K*_1_ and *K*_2_ denote the multisets of k-mers extracted from *s*_1_ and *s*_2_.

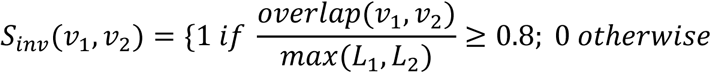

where *L*_1_ and *L*_2_ denote the inversion length of two haplotypes.

Two variant information from two haplotypes were grouped into a single diploid genotype (e.g., 1/1) if their computed similarity exceeded the similarity threshold of 0.8 (*--similarity_threshold*). This hierarchical approach ensures that the grouped variants share both high positional accuracy and high sequence identity.

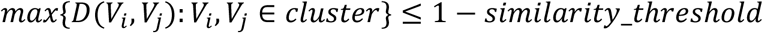

### DBSCAN clustering parameters

To evaluate the effect of DBSCAN clustering parameters on LGvar performance, we performed a parameter-sensitivity analysis using both human-human and human-chimpanzee genome comparisons. DBSCAN clustering distance parameters were set to 50 kbp, 100 kbp, 200 kbp, 500 kbp, and 1 Mbp. For the human–human comparison, SV calls from T2T-HG002 versus T2T-CHM13 were benchmarked against the GIAB HG002 CHM13v2.0-based benchmark VCF. For the human-chimpanzee comparison, where no gold-standard benchmark is available, we generated a high-confidence comparison set consisting of variants supported by at least two callers among PBSV, Sniffles, LGvar, PAV, SVIM-asm, and SyRI. For SDR benchmarking, we intersected the SDR coordinates identified by LGvar under different clustering parameters with genomic regions where alignments could not be uniquely assigned to a single genomic location.

### Assessment of assembly quality effects on SV detection

To assess the effects of sequencing depth and assembly quality on SV detection, HiFi and ONT reads from HG002 and chimpanzee were downsampled to 20×, 40× and 60× using rasusa^76^ (v4.1.0). The downsampled reads were assembled using hifiasm^77^ (v0.25.0-r726) with HiFi, ONT and Hi-C data. Assembly quality was assessed using Merqury^78^ (v1.3), with consensus QV calculated from k-mer spectra.

SVs were called from each assembly using LGvar, SVIM-asm and SyRI. For HG002, calls were benchmarked against the CHM13v2.0 GIAB HG002 SV benchmark. For the human– chimpanzee comparison, a consensus benchmark was constructed by merging SV calls from two read-based callers (PBSV and Sniffles) and four assembly-based callers (LGvar, PAV, SVIM-asm and SyRI) using SURVIVOR^79^. Precision, recall and F1 score were calculated for each sequencing-depth condition. Inversions were evaluated separately using the corresponding inversion benchmark sets.

LGvar accepts either genome assemblies or pre-computed alignment files as input. When assemblies are supplied, whole-genome alignments are generated internally with minimap2 (-cx asm20 --eqx --secondary=no -K 8G -t 12). Users who have already performed the alignment separately can supply the resulting PAF file directly, in which case LGvar bypasses the alignment step and produces identical SV calls.

### Benchmarking using simulated data

To evaluate the performance of LGvar relative to other assembly-based SV callers— including PAV^32^ (v2.4.6), SyRI^38^ (v1.7.0), and SVIM-asm^34^ (v1.0.3)—we generated simulated benchmark datasets using the VISOR^47^ framework (v1.1.2).

SVs were first randomly generated using the randomregion.r script provided by VISOR. In total, 500 SVs were simulated with the parameters -n 500 -l 10000 -s 5000 -v, encompassing deletions, insertions, inversions, tandem duplications, and cut-and-paste translocations at a predefined ratio of 30:30:30:5:5, respectively. To introduce realistic sequence divergence, high-confidence SNVs from the 1000 Genomes Project (GRCh38, phase 3 biallelic SNVs) were incorporated by adjusting the variant length and spacer parameters.

The simulated SVs and SNVs were integrated into chromosome 1 of the GRCh38 reference genome using the VISOR HACK module, generating a simulated haplotype assembly. Each SV caller was then applied using the simulated haplotype as the query assembly and the original GRCh38 chromosome 1 as the reference. Variant calls produced by each method were intersected with the ground-truth SV set to compute precision, recall, and F1 scores.

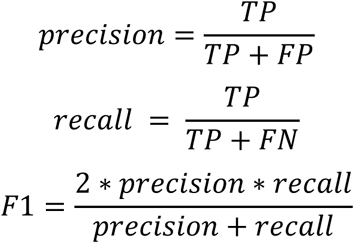

### Benchmarking in human population datasets

To assess LGvar performance on high-quality human genome assemblies, we benchmarked SV detection using validated datasets from the T2T-HG002 Q100 project and the Platinum Pedigree Consortium, including seven individuals (NA12877, NA12878, NA12879, NA12881, NA12882, NA12885, and NA12886) (https://github.com/Platinum-Pedigree-Consortium/Platinum-Pedigree-Datasets).

SVs were called using LGvar and three comparator assembly-based tools with default parameters. For SyRI, diploid variant call files were generated by merging haplotype-specific VCFs using bcftools^80^ (v1.20) merge to ensure compatibility with downstream benchmarking. SV calling performance was evaluated against curated truth sets using the bench module of Truvari^81^ (v5.4.0) with parameters -r 2000 -C 5000 --passonly --pick single --sizemax -1. SNVs and INDELs were benchmarked using hap.py with default settings. All benchmarking analyses were restricted to high-confidence genomic regions using the --includebed option.

To specifically evaluate inversion detection accuracy, we conducted a targeted benchmark across five human genomes (HG002, HG00733, HG02818, HG03486, and NA19240) provided in the GitHub (https://github.com/jamesc99/INV-Benchmark/tree/main/data_zenodo/bed). Unique inversion calls from each method were manually inspected using dot plots generated with MUMmerplot^82^ (v3.5) to validate breakpoint structure and orientation.

### Benchmarking in cross-species datasets

#### Human-macaque large-scale genomic differences

To evaluate the robustness of LGvar in cross-species analyses, we applied LGvar together with the two comparator tools to a genome-wide comparison between the human ( T2T-CHM13) assembly and the crab-eating macaque assembly (*Macaca fascicularis*, T2T-MFA8).

To ensure coordinate consistency between assemblies, chromosomes were reverse-complemented when necessary based on alignment orientation. Specifically, a chromosome was reverse-complemented if (i) the cumulative length of inverted alignments exceeded 50% of the chromosome length, or (ii) both terminal regions exhibited inverted orientations. Based on these criteria, T2T-CHM13 chromosomes 1, 3, 6, 9, 18, 21, and 22, as well as T2T-MFA8 chromosome 13, were reverse-complemented prior to downstream analysis.

Whole-genome pairwise alignment was performed using minimap2 (v2.30-r1287) with T2T-MFA8 as the reference and T2T-CHM13 as the query sequence. Alignment was run with the following parameters: -a -x asm20 --eqx --cs -K 500M -k 15 -m 10 -A 1 -B 2 -O 2,12 -n 2 -g 100 -r 200,100000 --secondary=no -s 1000 -o aln.sam. The resulting SAM-format alignments were used as input for SyRI and SVIM-asm. For LGvar analysis, SAM alignments were converted to PAF format using paftools.js sam2paf (minimap2 package). LGvar was executed with parameters -m sensitive -dv 0.2 -d 2000.

We used a curated set of large-scale genomic rearrangements reported by Zhang *et al.* as the benchmark reference. Overlap between tool-derived variant calls and the curated events was assessed using bedtools^83^ (v2.31.1) intersect with parameters -f 0.5 -F 0.5, requiring at least 50% reciprocal overlap to define a true positive. Owing to computational constraints, SV calling with PAV was performed on a per-chromosome basis, whereas LGvar and SVIM-asm were run genome-wide in a single execution.

#### Plant genome comparison

To further assess LGvar performance in plant genomes, we analyzed SVs between two *Arabidopsis thaliana* assemblies, using TAIR10 (GCA_000001735.3) as the reference and the Ler accession (GCA_900660825.1) as the query. SyRI has previously been applied to this genome pair and reported its genomic rearrangement events, which we used as a control.

We applied LGvar to the same dataset and performed parallel analyses using SVIM-asm and PAV. Using criteria consistent with those described above, we compared the variant calls across all tools. To assess accuracy, we randomly selected 100 insertions, 100 deletions, and 100 inversions per tool (300 events per caller) for manual inspection.

LGvar exhibited the lowest FP rates (12.08%) among all methods and identified nearly 1.5-fold as many high-confidence events as SyRI, demonstrating improved sensitivity while maintaining high precision in plant genome comparisons.

### Large-scale lineage-specific fixed SVs

#### Lineage-specific SVs callset

To assess the generalizability of LGvar, we identified insertions, deletions, and inversions across two human genomes (HG002 and C088) and five NHP genomes (chimpanzee, bonobo, gorilla, Bornean orangutan, and Sumatran orangutan), using the T2T-CHM13 assembly as the reference. Structural variants (SVs) detected by each caller in each sample were merged using SV-pop^32^ to generate a unified NHP SV callset. SVs located within complex genomic regions were excluded, and only homozygous SVs of at least 10 kbp were retained and considered fixed within each lineage. To reduce potential reference bias, we performed a reciprocal analysis in which the Sumatran orangutan genome was used as the reference, and all remaining genomes were treated as queries.

To further minimize technology- and algorithm-specific detection biases, candidate SVs were subjected to rigorous manual curation. Alignment visualizations were generated using MUMmerplot, SafFire^84^ (https://github.com/mrvollger/SafFire), and IGV^85^ (v2.19.4) to resolve ambiguous or complex regions. Each candidate SV was systematically inspected and manually classified as a TP or FP based on hands-on checks from the alignments.

#### Reference bias analysis

Because genome alignment is inherently reference dependent, reference bias may influence SV discovery. To assess the extent of reference bias in our analyses, we performed reciprocal SV calling using T2T-CHM13 and T2T-Sumatran orangutan assemblies as references, as described above.

SV callsets generated from the two reference-based analyses were compared by lifting over the genomic coordinates of large-scale insertions and deletions between the two reference genomes using transanno (https://github.com/informationsea/transanno, v0.4.5). Inconsistent calls were further manually inspected using MUMmerplot and SafFire. We identified 817 fixed insertions and deletions that were consistently detected in both reference-based callsets, whereas 728 and 483 events were uniquely detected when using the T2T-CHM13 and Sumatran orangutan references, respectively. These results indicate that reference bias affects 59.7% of large-scale insertion and deletion calls (728+483) / (728+483+817), leading to inconsistent detection across references.

In contrast to insertions and deletions, inversions are difficult to reliably lift over between different reference genomes. Therefore, inversion curation was performed on a chromosome-by-chromosome pair comparison by hand. Unique inversion calls from each SV callset were manually inspected, resulting in a final set of 174 fixed inversions.

Next, we classified SVs of at least 10 kbp as lineage-specific fixed events. In this framework, human-specific deletions were defined as loci present in non-human primate genomes but absent from the human lineage, whereas Pan-specific insertions were defined as sequences present in chimpanzee and bonobo genomes but absent from the human and other primate genomes. This classification enabled reconstruction of lineage-specific evolutionary SV events. Finally, we calculated the large-scale SV-to-SNV ratio following the same approach as described in previous studies.

#### Human and nonhuman great ape inversion

Previous studies have reported 338 lineage-specific inversions across the great ape clade^40^. In this study, we restricted our analysis to large-scale and fixed inversions that do not overlap centromeric regions, yielding a curated set of 127 inversions.

An inversion in our dataset was considered a true positive if it showed a reciprocal overlap greater than 0.5 with a previously reported event and was assigned to the same evolutionary lineage. Under these criteria, LGvar successfully recapitulated 91.3% (116/127) of the benchmarked inversions. The remaining eleven events were classified as syntenic in our analysis, likely reflecting differences in genome assembly quality and contiguity between the non-human great ape assemblies used here and those employed in earlier studies.

#### Genes disrupted by large-scale lineage-specific fixed SVs

To ensure non-redundant annotation, we identified representative transcripts by prioritizing ‘RefSeq Select’ entries; where unavailable, the longest transcript was selected for subsequent intersection analysis.

For lineage-specific deletions, SV coordinates were intersected with gene annotations from the reference genome. For lineage-specific insertions, the coordinates of query genomes were extracted from the VCF files produced by LGvar and PAV, then the coordinates were intersected with gene annotations from the corresponding query genome, using the primate genome annotations provided by the MARBL lab (https://github.com/marbl/Primates). For lineage-specific inversions, gene disruption was assessed by intersecting inversion breakpoints with the T2T-CHM13 genome annotation.

Using this approach, we identified 59 protein-coding genes that were entirely gained through insertions and 66 genes that were completely lost through deletions. In addition, 9 and 14 protein-coding genes were partially disrupted within exonic regions by insertions and deletions, respectively. Inversion events also contributed to gene disruption, with inversion breakpoints directly intersecting exonic regions of an additional 4 protein-coding genes.

#### Genes disrupted by large-scale SDRs

To identify comprehensive SDR regions for the five NHP genomes (chimpanzee, bonobo, gorilla, Bornean orangutan, and Sumatran orangutan), we extracted SDR calls from LGvar by two mapping directions and incorporated the centromeric and telomeric regions that were previously masked.

These SDR coordinates were intersected with their primary gene annotations to extract protein-coding genes. For protein-coding genes in each species, only the transcript with the longest CDS was retained. These representative sequences were then used to construct a custom database via makeblastdb. We performed an all-vs-all alignment for CDS of these SDR region genes via blastn^86^ (v2.17.0) with parameters *-outfmt “6 qseqid sseqid pident length qlen slen qstart qend sstart send evalue bitscore” -evalue 1e-10 -qcov_hsp_perc 50 - max_target_seqs 100*. To ensure unique records, alignments were filtered by gene name and identity.

Copy number analysis identified 659 gene gain/loss events within SDR regions. Finally, disrupted genes by SVs and SDRs were benchmarked against datasets from Mao *et al.* (Cell, 2024)^12^ and Sudmant *et al.* (Genome Res, 2013)^55^.

Then we performed Gene Ontology (GO) enrichment analysis of the entire genes affected by SVs and SDRs using the Database for Annotation, Visualization, and Integrated Discovery (DAVID)^87^.

## Authors’ contributions

Y.M. conceived the project. F.Z., J.H., and S.Z. developed and validated the tool. F.Z. and J.H. performed all data analysis. J.L. and E.E.E. provided the great ape genomes and benchmarking datasets. F.Z. and Y.M. drafted the manuscript. All authors read and approved the manuscript.

## Data Availability

The code for LGvar is available on GitHub (https://github.com/YafeiMaoLab/LGvar) with an MIT license. The assemblies or genomic sequences used in this study can be found in GenBank with the accession numbers GCA_009914755.4, GCA_018852605.3, GCA_000001405.15, GCA_028858775.2, GCA_028885625.2, GCA_028885655.2, GCA_029281585.2, GCA_029289425.2, GCA_000001735.3, GCA_037993035.2, and GCA_900660825 and the github link (https://github.com/Platinum-Pedigree-Consortium/Platinum-Pedigree-Datasets).

